# Rab27b regulates the release, autophagic clearance, and toxicity of alpha-synuclein

**DOI:** 10.1101/2020.03.09.984096

**Authors:** Rachel Underwood, Bing Wang, Christine Carico, Robert H. Whitaker, William J. Placzek, Talene Yacoubian

## Abstract

Alpha synuclein (αsyn) is the primary component of proteinaceous aggregates termed Lewy Bodies that pathologically define synucleinopathies including Parkinson’s disease (PD) and Dementia with Lewy Bodies (DLB). αSyn is hypothesized to spread through the brain in a prion-like fashion by misfolded protein forming a template for aggregation of endogenous αsyn. The release and uptake of αsyn from cell to cell are considered important processes for this prion-like spread. Rab27b is one of several GTPases essential to the endosomal-lysosomal pathway and is implicated in protein secretion and clearance but has yet to be characterized in its role in αsyn spread. In this study, we used a paracrine αsyn *in vitro* model to test the impact of Rab27b on αsyn release, clearance, and toxicity. shRNA-mediated knockdown (KD) of Rab27b increased αsyn-mediated paracrine toxicity. While Rab27b reduced αsyn release primarily through non-exosomal pathways, the αsyn released under KD conditions was of higher molecular weight species by size exclusion chromatography. Rab27b KD increased intracellular insoluble αsyn levels and led to an accumulation of endogenous LC3 positive puncta. Rab27b KD also decreased LC3 turnover with chloroquine treatment, indicating a defect in autophagic flux. Rab27b protein levels were increased in postmortem human brain lysates from PD and DLB subjects compared to healthy controls. These data indicate a role for Rab27b in the release, clearance, and toxicity of αsyn and ultimately in the pathogenesis of synucleinopathies.

## INTRODUCTION

Synucleinopathies, such as Parkinson’s Disease (PD) and Dementia with Lewy Bodies (DLB), have tremendous economic and social impact, yet there are no current therapies to slow neurodegeneration in PD or DLB (1). Alpha-synuclein (αsyn) is the key pathogenic protein implicated in both PD and DLB. Recent evidence suggests that misfolded αsyn propagates from neuron to neuron in a prion-like manner: transmission of misfolded αsyn templates misfolding of endogenous αsyn to promote further aggregation and ultimately neurotoxicity (2-4). Propagation of αsyn requires three distinct cellular processes: release, uptake, and misfolding. αSyn does not have a signal peptide and is not released via the classic ER-Golgi secretory pathway (5,6). There is evidence that αsyn is released through non-classical endosomal pathways, including exosomal, misfolding-associated protein secretion, and lysosomal pathways (7-13). These pathways are thought to be tightly regulated by the Rab family of GTPase proteins.

Rab GTPases are a largely conserved family of proteins with over 60 known mammalian members. Rab proteins’ primary functions are carried out through a catalytic GTP/GDP binding site which, when GTP bound, causes a conformational switch allowing the protein to interact with its effector proteins, and thus regulate multiple cellular processes (14). Several Rab GTPases have been associated with PD. Mutations in Rab39b and Rab32 cause autosomal recessive forms of early onset and late onset PD, respectively (15-17). Rabs regulate autophagic, exocytotic, and endocytotic pathways potentially involved in αsyn transmission. A number of Rab GTPases regulate αsyn trafficking (18). Overexpression of Rab1, 7, 8, or 11 is protective against αsyn toxicity, and these Rabs are implicated in αsyn aggregate formation in PD models (19-24). Additionally, Rabs have been shown to be kinase substrates of LRRK2, mutations of which are the most common genetic cause of PD (25).

Rab27b is a relatively unstudied member of the Rab GTPase family. Pointing to a potential functional overlap between the two proteins, Rab27b shares common regulators and effectors with Rab3, which binds αsyn and is localized in αsyn aggregates (14,22,26-28). Rab27b can regulate protein secretion through both exosomal and non-exosomal pathways by regulating transport and docking steps in tandem with Rab27a through interaction with several Rab27 effectors (29-35). Rab27b and its effectors regulate exocytosis of dense-core vesicles in neuronal lines and synaptic vesicle release in neurons (27,28,36-38). Unlike Rab27a, Rab27b is highly expressed in neurons in the cortex, striatum, and midbrain, areas affected in PD and DLB (29,39). Rab27b may also play a role in protein clearance via autophagy. Rab27b is localized to the autophagosome under stress conditions that induce autophagy (40). Rab27a/b double knockout mice (DKO) show increased vesicle formation, including lysosomes and autophagosomes, in the lacrimal gland (41). Evidence for autophagic-lysosomal pathway impairment in PD is growing (42-46). The presence of Rab27b in the autophagic-lysosomal process indicates the potential for Rab27b to alter not only αsyn release but αsyn clearance as well.

Rab27b has been linked to several neurodegenerative disorders. Elevated Rab27b expression in cholinergic basal forebrain neurons is associated with cognitive decline in mild cognitive impairment and Alzheimer’s disease (47). In addition, multiple Rab27b polymorphisms have been associated with a higher risk for motor neuron disease in GWAS studies (48). Alterations in Rab27b expression are observed in a cellular model for X-linked Dystonia Parkinsonism syndrome (XDP) and in human DLB brains (49,50).

Because of its function in regulating protein secretion and autophagy, we hypothesized that Rab27b regulates cell to cell transmission of αsyn by regulation of αsyn release and clearance. Here we examine the effect of Rab27b on αsyn toxicity in an *in vitro* paracrine αsyn model, and evaluate the impact of Rab27b on αsyn release and clearance via autophagy. We found that Rab27b reduces αsyn toxicity by promoting autophagic flux.

## RESULTS

### Rab27b reduces αsyn induced paracrine toxicity

We have developed a paracrine αsyn model using a doxycycline (doxy)-inducible neuroblastoma cell line, termed isyn, in order to evaluate the toxicity associated with neuronally-released αsyn (Fig. 1a) (51). We have previously shown that induction of αsyn expression with doxy in isyn cells leads to the detection of αsyn in the CM in a dose-dependent manner, and that fractionation of the CM into exosomal and non-exosomal fractions by sequential, high ultracentrifugation techniques shows that αsyn is released into both exosomal and non-exosomal fractions (51). Additionally, we have previously established that released αsyn is toxic to separately cultured neurons: transfer of αsyn-enriched CM from induced isyn cells promotes cell death of separately cultured differentiated SH-SY5Y neuroblastoma cells or primary mouse neurons (51). Toxicity from αsyn-enriched CM depends on αsyn, as immunodepletion of αsyn from the CM eliminated toxicity in a dose-dependent manner (51). An initial PCR screen for detectable expression of potential exosome-related proteins in isyn cells pointed to Rab27b and Rab35 expression in isyn cells. Upon αsyn induction with doxy (10μg/ml), Rab27b expression increased by nearly 2-fold, as determined by immunocytochemistry (Fig. 1b). We confirmed the increase in Rab27b by Western blot (Supp. Fig. 1). No change was noted in Rab35 expression (Supp. Fig. 2a).

**Figure 1.**
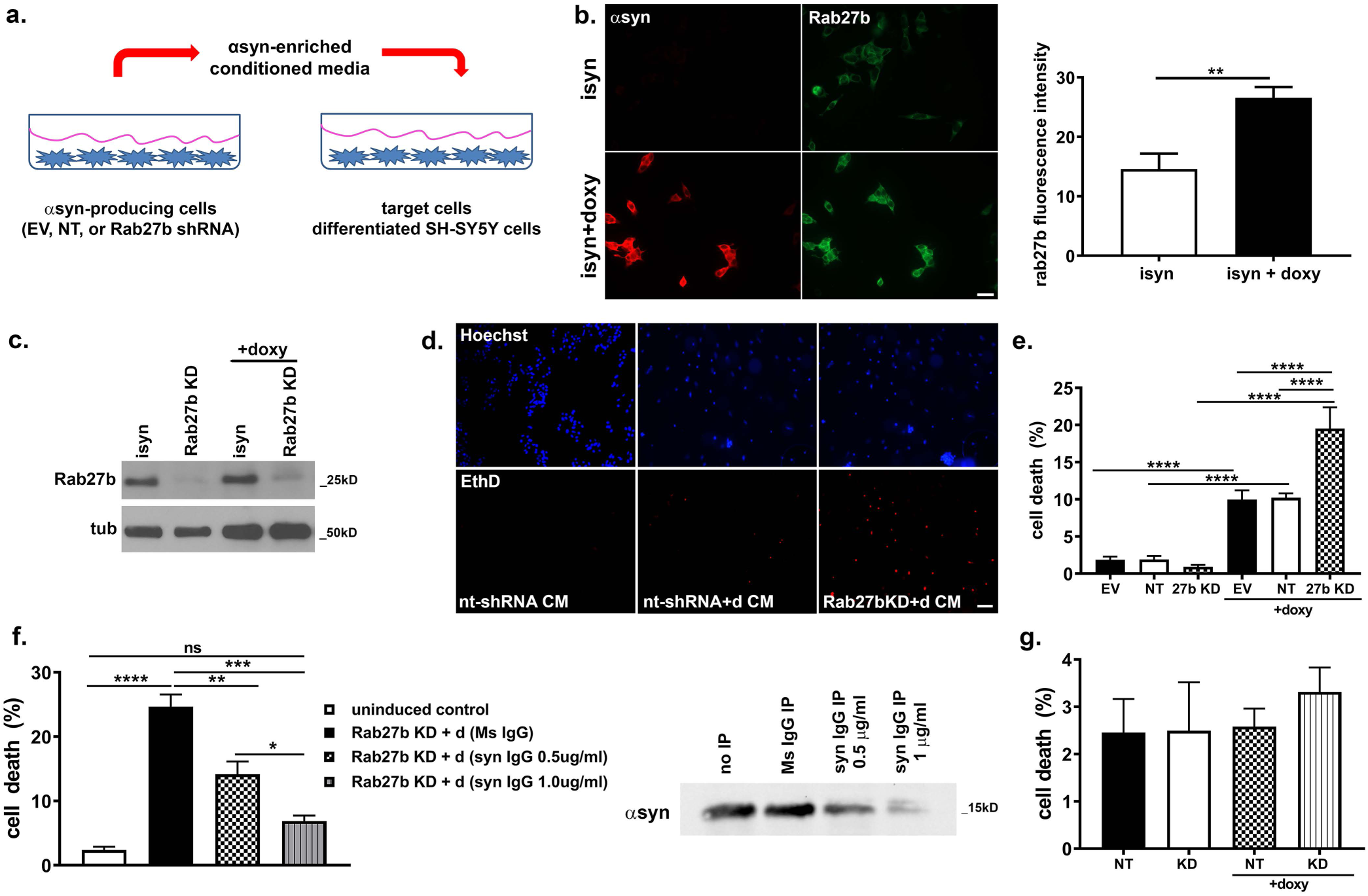
Rab27b knockdown in isyn cells increases αsyn-induced paracrine toxicity. a) Schematic of αsyn paracrine toxicity model. Isyn cell transfected with plko.1 empty vector (EV), non-targeted shRNA (nt-shRNA), or Rab27b shRNA overexpress and release αsyn into conditioned media (CM) when treated with doxy. Cell death is induced in recipient cells treated with αsyn-enriched CM. b) Rab27b immunoreactivity increased in isyn cells upon αsyn overexpression. Representative images of Rab27b (green) and αsyn (red) immunoreactivity in uninduced isyn cells and in isyn cells when induced with doxy for 48h. Scale bar = 25 μm. n = 4. Student t-test t_6_ = 3.793, **p≤0.01. c) Rab27b KD by shRNA reduces Rab27b protein levels in both uninduced and induced isyn cells. d) Rab27b KD increases αsyn-induced paracrine toxicity in differentiated SH-SY5Y cells treated with CM. Representative images of differentiated SH-SY5Y cells treated with CM from isyn cells or isyn cells transduced with non-target (nt) or Rab27b shRNA. Ethidium D labels the nuclei of dying cells, while Hoechst 33342 stains the nuclei of all cells. Scale bar = 50 μm. e) Quantification of cell death in differentiated SH-SY5Y cells treated with CM from isyn cells transduced with empty vector (EV), non-target (nt) shRNA, or Rab27b shRNA for 48 hours. n = 3 independent rounds with 1-2 replicates per round. One-way ANOVA: F_(3, 8)_ = 44.90, p≤0.0001. Tukey’s multiple comparison test: ****p≤0.0001. f) Immunoprecipitation of αsyn from CM by αsyn-directed monoclonal antibody reduces the toxicity of αsyn-enriched CM from isyn/Rab27b KD cells in a dose-dependent manner. Quantification of cell death in differentiated SH-SH5Y cells treated with CM from induced isyn/Rab27b KD cells in which αsyn was immunodepleted. Western blot of CM after immunoprecipitation confirms reduction of αsyn in the CM. n = 3 independent rounds with 1 replicate per round. One-way ANOVA: F_(5, 20)_ = 39.10, p≤0.0001. Tukey’s multiple comparison test: *p≤0.05, **p≤0.01, ***p≤0.001, ****p≤0.0001. g) Rab27b KD does not induce cell death in isyn cells. Quantification of cell death in isyn transduced with nt-shRNA or Rab27b shRNA with and without doxy induction. Number of Ethidium D positive cells is normalized to total cell count, as determined by Hoechst 33342 staining. n = 3 independent rounds with 1 replicate per round. One-way ANOVA: F_(3, 8)_ = 0.3343, p>0.05. Error bars represent SEM. D = doxy.

Given the increase in Rab27b expression upon αsyn induction in isyn cells, we tested whether Rab27b knockdown (KD) could affect the toxicity of released αsyn. Isyn cells were transduced with Rab27b targeted shRNA lentivirus selectable by puromycin. Rab27b KD in isyn cells was confirmed by Western blot (Fig. 1c). As controls for Rab27b KD, isyn cells were transduced with either plko.1 empty vector (EV) lentivirus or with nontarget shRNA (nt-shRNA) lentivirus. Toxicity of αsyn-enriched CM was increased with Rab27b KD upon doxy induction in isyn cells compared to both isyn controls. Separately-cultured differentiated SH-SY5Y showed ∼10% cell death at 24 hours when treated with CM from induced control isyn cells, but toxicity was doubled when treated with CM from induced isyn cells with Rab27b KD (Fig. 1d, e). Increased toxicity from αsyn-enriched media from Rab27b KD/isyn cells was dependent on αsyn, as immunodepletion of αsyn from the CM eliminated toxicity in a dose dependent manner (Fig. 1f). Rab27b KD did not impact cell death at baseline in isyn cells (Fig. 1g).

### Rab27b regulates αsyn release

Rab27b regulates protein secretion through both exosomal and non-exosomal pathways (29-35). As Rab27b KD increased the toxicity of αsyn released by isyn cells, we next examined whether Rab27b regulated the amount of αsyn released in this paracrine model system. Surprisingly, upon doxy induction we observed a nearly 60% decrease in the total amount of αsyn released into the CM in isyn cells with Rab27b KD compared to control isyn cells transduced with nt-shRNA (Fig. 2). Differences in release were not due to cell death, as cell death was limited to ∼2-4% in induced isyn cells with and without Rab27b KD (Fig. 1g). These data indicate that the increase in αsyn toxicity with Rab27b KD was not due to a total increase in αsyn release.

**Figure 2.**
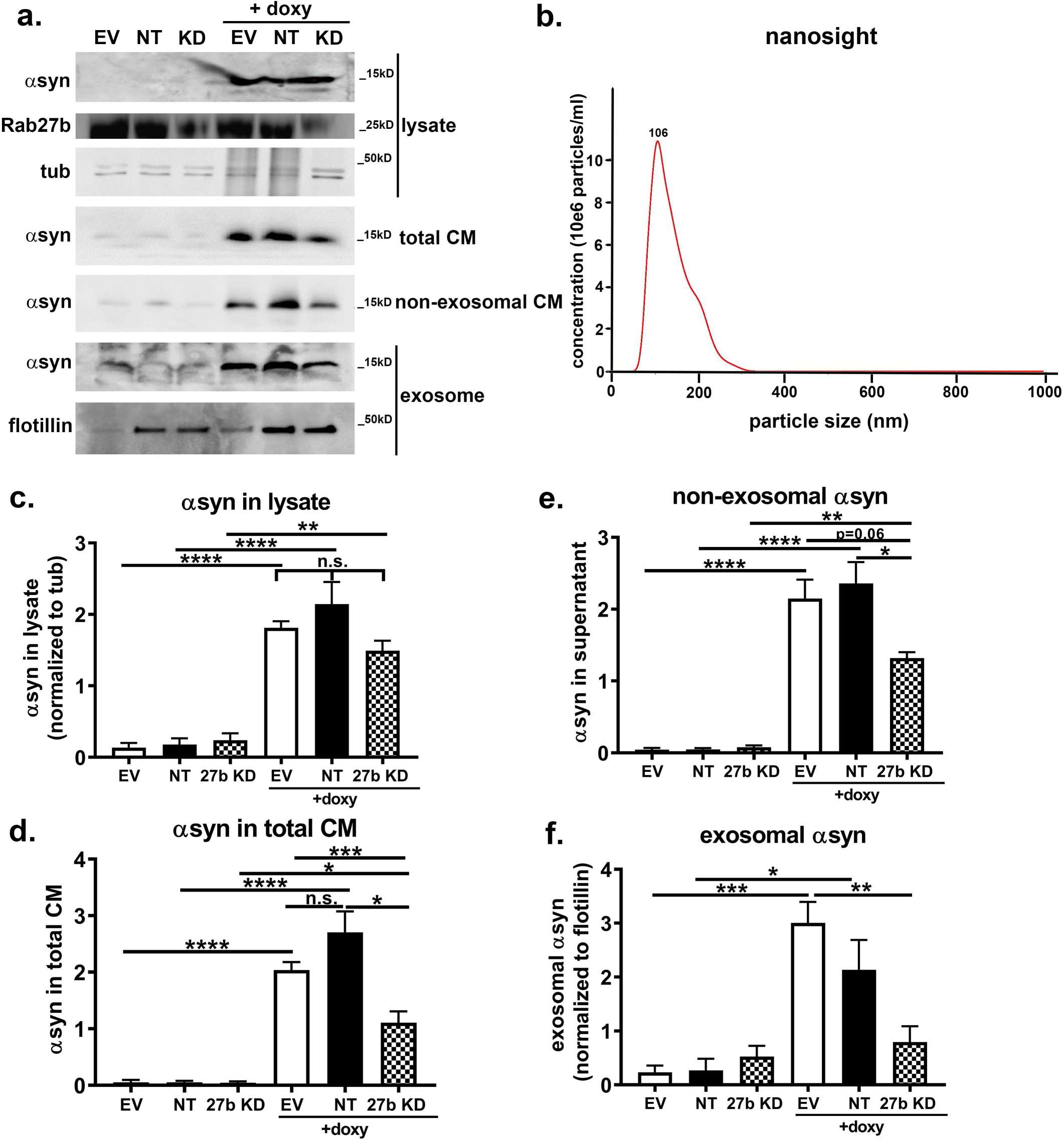
Rab27b knockdown reduces αsyn release into the CM. a) Representative Western blots of total αsyn levels in CM and lysates from uninduced and induced isyn cells lines transduced with empty vector (EV), nt-shRNA, or Rab27b shRNA after doxy (10 μg/ml) induction for 96 hours. Equal protein levels were loaded for each CM sample. b) Nanosight shows that exosomal fraction from CM of control nt-shRNA transduced isyn cells contains particles averaging 106 nm, consistent with the size of exosomes, after 96 hours of induction with doxy. c) Quantification of αsyn in the intracellular lysates from isyn cells lines transduced with EV, nt-shRNA, or Rab27b shRNA with and without 96 hour induction by Western blot. αSyn in lysates was normalized to tubulin. n = 3 independent rounds. One-way ANOVA: F_(5, 12)_ = 34.43, p≤0.0001. Sidak’s multiple comparison test: **p≤0.01, ****p≤0.0001. n.s. = non-significant. d) Quantification of αsyn in total, unfractionated CM from isyn cells lines transduced with EV, nt-shRNA, or Rab27b shRNA with and without 96 hour induction by Western blot. Equal protein amounts were loaded for each CM sample. n = 3 independent rounds. One-way ANOVA: F_(5, 12)_ = 40.21, p≤0.0001. Sidak’s multiple comparison test: *p≤0.05, ***p≤0.001, ****p≤0.0001. n.s. = non-significant. e) Quantification of αsyn in the non-exosomal fraction from the CM from isyn cells lines transduced with EV, nt-shRNA, or Rab27b shRNA with and without 96 hour induction by Western blot. Equal protein amounts were loaded for each non-exosomal fraction sample. n = 3 independent rounds. One-way ANOVA: F_(5, 12)_ = 43.13, p≤0.0001. Sidak’s multiple comparison test: *p≤0.05, ***p≤0.001, ****p≤0.0001. f) Quantification of αsyn in the exosomal fraction from CM from isyn cells lines transduced with EV, nt-shRNA, or Rab27b shRNA with and without 96 hour induction by Western blot. Exosomal αsyn was normalized to flotillin. n = 3 independent rounds. One-way ANOVA: F_(5, 12)_ = 12.20, p≤0.001. Sidak’s multiple comparison test: *p≤0.05, ***p≤0.001, ***p≤0.001. Error bars represent SEM. D = doxy.

Previous research has shown that αsyn is released through exosomal and non-exosomal pathways, and certain studies have suggested that exosomally-released αsyn may have increased toxic potential (5,11,52,53). We fractionated the CM into exosomal and non-exosomal fractions by sequential, high ultracentrifugation techniques (54) to test if Rab27b KD possibly preferentially inhibited release through non-exosomal pathways. The vast majority of released αsyn in our model is associated with the non-exosomal fraction from isyn cells (51). When Rab27b was knocked down in isyn cells, the amount of αsyn released in the non-exosomal fraction was decreased by 44% compared to nt-shRNA control (Fig. 2e). The amount of αsyn released in the exosomal fraction was also decreased relative to control isyn cells (Fig. 2f). Nanosight analysis confirmed nanometer-sized vesicle sizes consistent with exosomes released into the CM (51) (Fig. 2b). Thus, the total change in αsyn release induced by Rab27b KD occurred through non-exosomal and exosomal pathways.

### Rab27b promotes autophagic flux and αsyn clearance

Our previous work in the paracrine αsyn model revealed that the type of αsyn species released was the critical factor that mediates toxicity, not the total amount of αsyn released (51). This finding suggests that oligomeric, toxic αsyn species are potentially available for release. To biochemically characterize αsyn released into the CM, we used size exclusion chromatography (SEC). We and others have previously shown that much of the released αsyn is found in higher molecular weight fractions representing molecular sizes greater than 14kD, the expected monomeric αsyn size by SEC (5,51,55). We fractionated CM from induced nt-shRNA/isyn cells and induced isyn/Rab27b KD cells by SEC and found that the amount of released αsyn released into higher molecular weight fractions was significantly increased by Rab27b KD (Fig. 3a, b). ELISA of total CM prior to SEC fractionation confirmed that total αsyn levels was overall lower in the CM from isyn cells with Rab27b KD compared to that from induced nt-shRNA isyn cells despite the increase in higher molecular weight αsyn.

**Figure 3.**
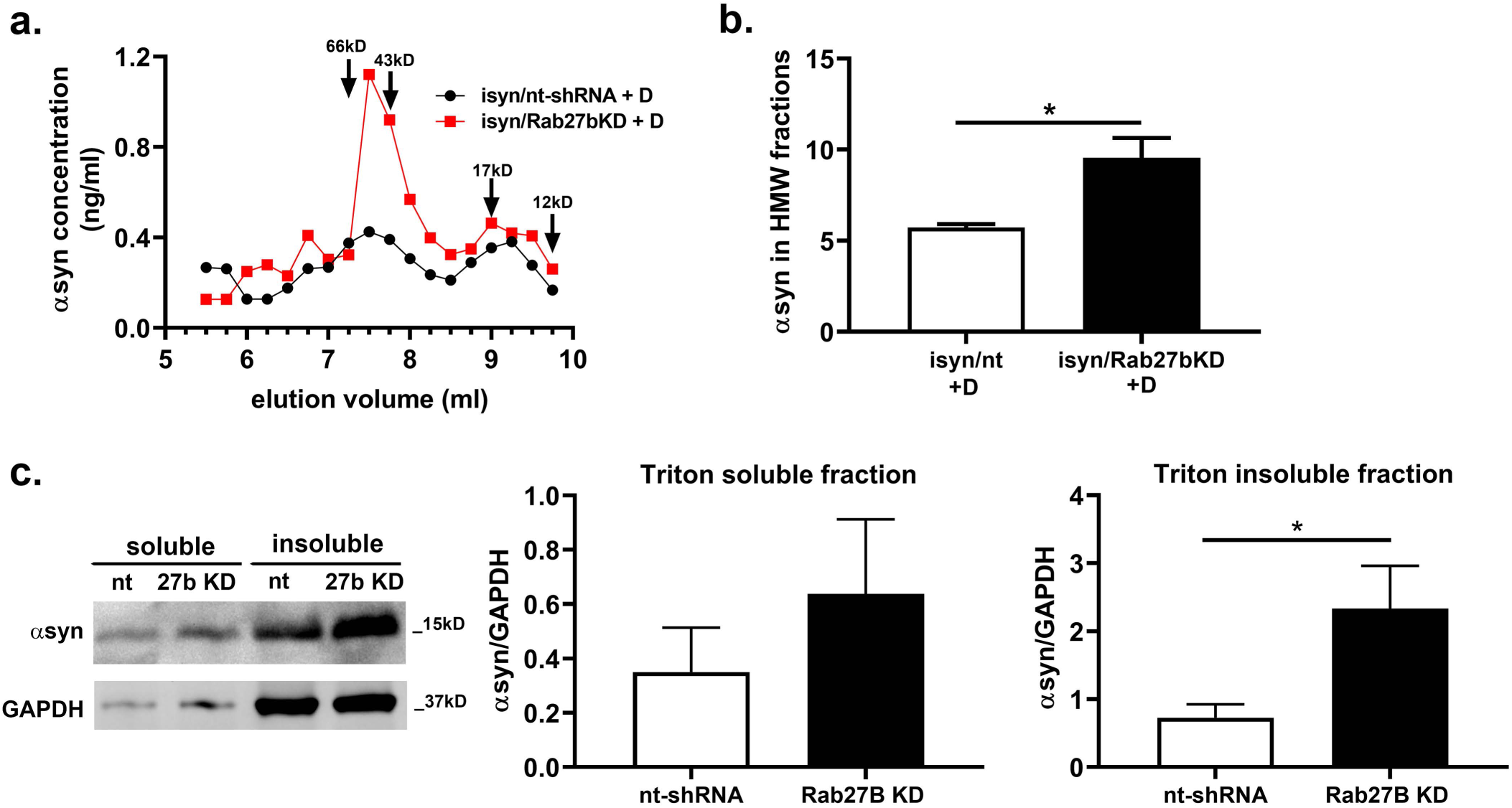
Rab27b knockdown increases release of higher molecular weight species of αsyn. a) αSyn levels in SEC fractions of CM from induced isyn/nt-shRNA cells or from induced isyn/Rab27b KD cells. Equal protein amounts (80μg) per sample were loaded onto the column, and 250 μl fractions were collected from elution volume 4 – 12.5 ml. αSyn in each fraction was measured by ELISA. αSyn from CM from induced isyn/Rab27b cells was partially shifted into higher molecular weight fractions. Data is representative of four independent experiments. b) Quantification of released αsyn detected in high molecular weight (HMW) SEC fractions collected between elution volumes 7.25 to 8.5 ml. n = 4 independent rounds. Student t-test t_6_ = 3.484, *p≤0.05 (Student’s t-test). c) Representative Western blot and quantification of αsyn in Triton X100 soluble and insoluble fractions from induced isyn cells lines transduced with nt-shRNA (nt) or Rab27b shRNA after doxy (10 μg/ml) induction for 96 hours. αSyn was normalized to GAPDH. n = 4 independent rounds. Student t-test t_6_ = 2.43, *p≤0.05 (Student’s t-test). Error bars represent SEM. D = doxy.

Given the increase in HMW αsyn species released in the presence of Rab27b KD, we hypothesized that Rab27b promotes the clearance of intracellular αsyn. We next examined whether insoluble αsyn levels were altered in induced isyn cells upon Rab27b KD. While αsyn levels in the Triton X-100 soluble fraction did not statistically differ, we observed more αsyn in the Triton X-100 insoluble fraction in isyn cells with Rab27b KD (Fig. 3c).

Misfolded proteins are typically cleared through the autophagic pathway or through the proteasome. αSyn, particularly oligomeric species, has been previously shown to be degraded through autophagy (46,56). We hypothesized that Rab27b promotes αsyn clearance via the autophagic-lysosomal pathway. We first examined levels of proteins involved in the endosomal-lysosomal system in doxy-induced isyn with and without Rab27b KD cells under serum starvation. Rab7, a late endosome marker, and p62, an autophagy marker were increased in Rab27b KD cells, while early endosome marker Rab5 remained unchanged (Fig. 4). Similarly, LC3-positive puncta were increased in induced isyn cells with Rab27b KD under serum starvation compared to induced control isyn cells (Fig. 5a).

**Figure 4.**
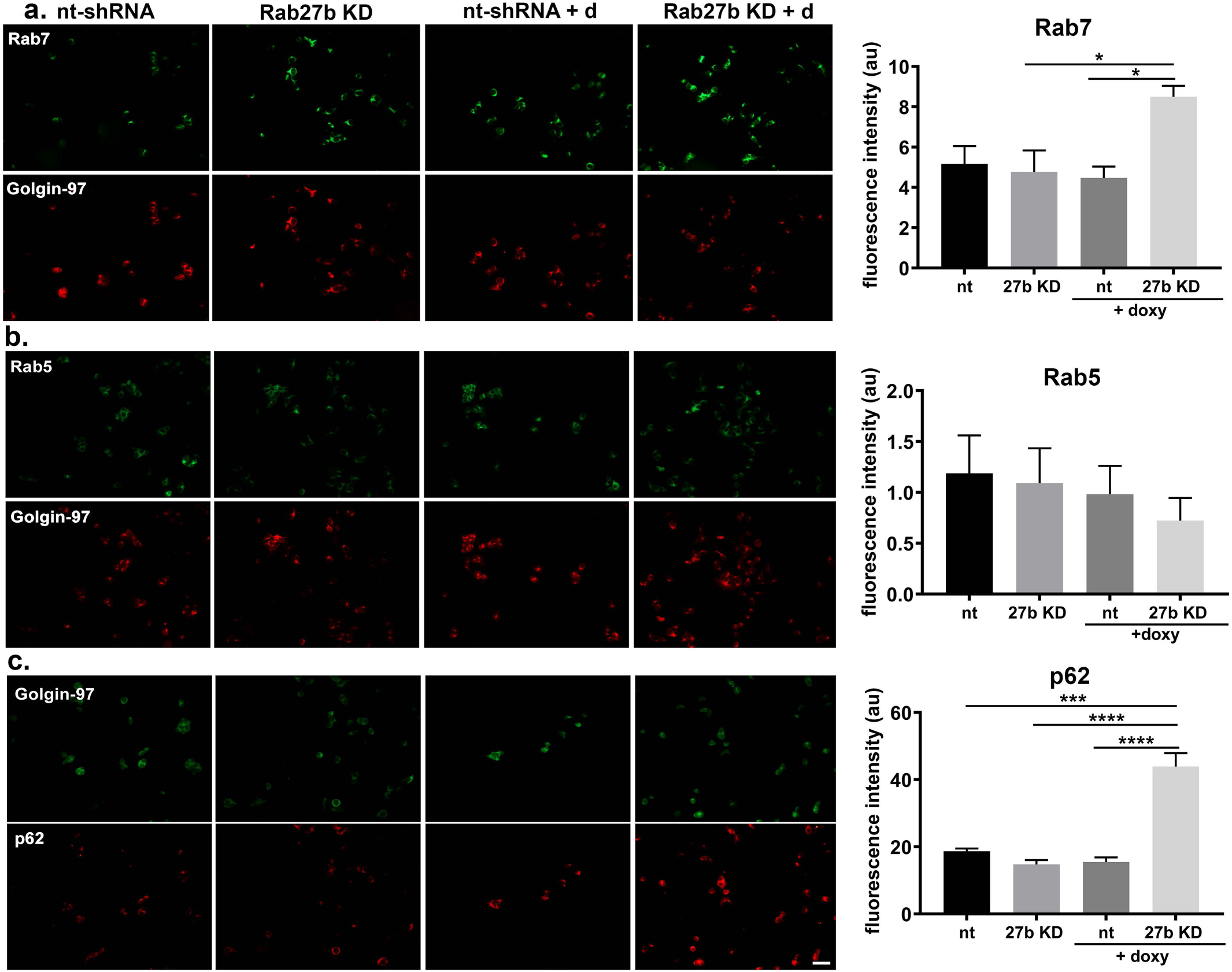
Rab27b knockdown increases Rab7 and p62 in isyn cells. a) Representative images and quantification of Rab7, a late endosomal marker, in isyn cells with and without Rab27b KD when induced with doxy for 48h. Rab7 fluorescence intensity is normalized to Golgin-97, a marker of early trans-Golgi network membranes. n = 3 independent rounds. One-way ANOVA: F_(3, 8)_ = 5.443, p≤0.05. Tukey’s multiple comparison test: *p≤0.05. b) Representative images and quantification of Rab5, an early endosomal marker, in isyn cells with and without Rab27b KD when induced with doxy for 48h. Rab5 fluorescence intensity is normalized to Golgin-97. n = 3 independent rounds with 1-2 replicates per round. One-way ANOVA: F_(3, 11)_ = 0.4304, p>0.05. c) Representative images and quantification of p62 in isyn cells with and without Rab27b KD when induced with doxy for 48h. p62 fluorescence intensity is normalized to Golgin-97. n = 3 independent rounds. One-way ANOVA: F_(3, 8)_ = 37.57, p≤0.0001. Tukey’s multiple comparison test: ***p≤0.001, ****p≤0.0001. Error bars represent SEM. D = doxy. Scale bar = 50 μm.

**Figure 5.**
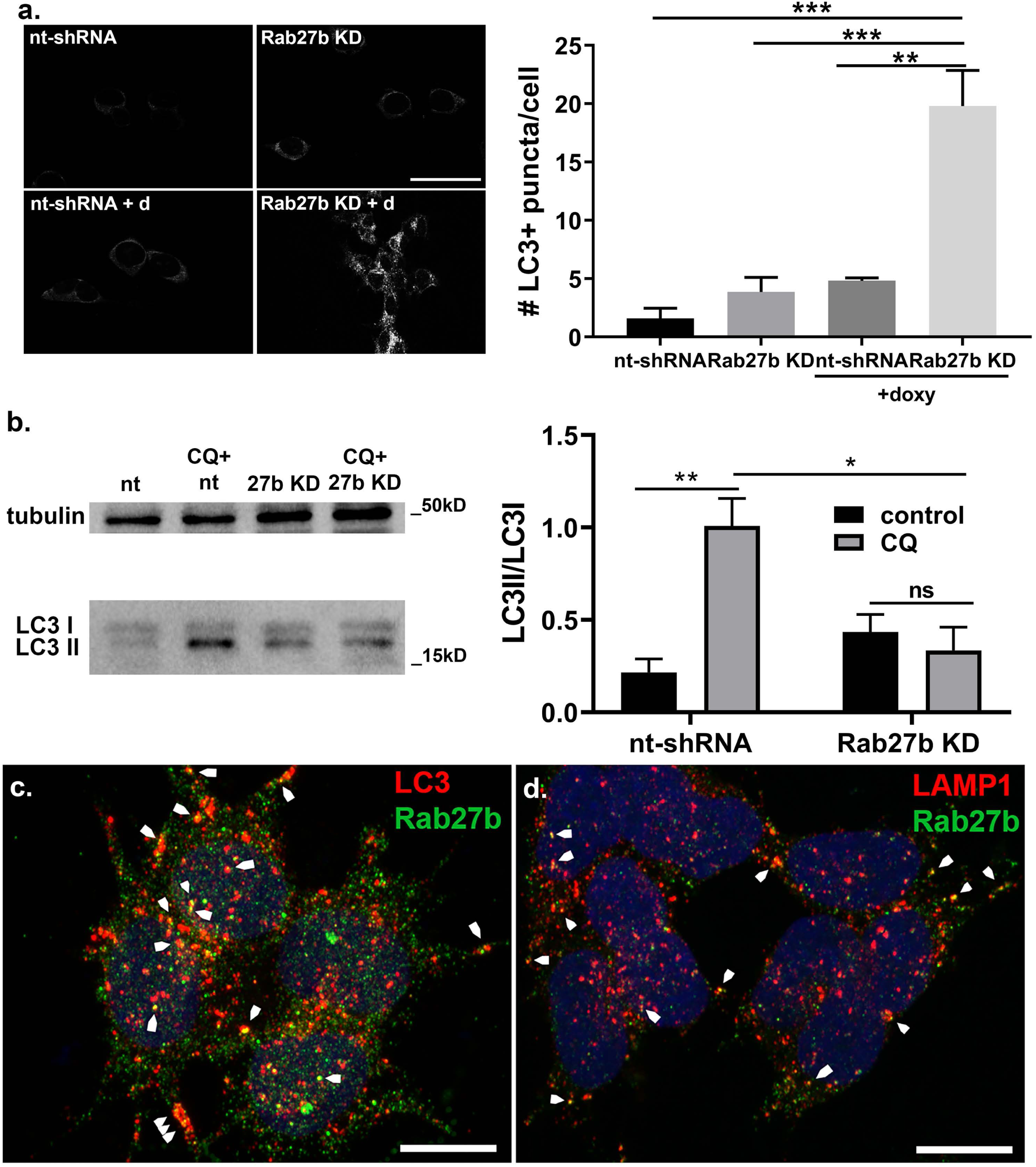
Rab27b knockdown reduces autophagic flux in isyn cells. a) Rab27b KD decreases LC3 puncta numbers in comparison to control. Representative images and quantification of LC3-positive puncta in isyn cells with and without Rab27b KD when induced with doxy for 48h. Scale bar = 50 μm. n = 3 independent rounds, 17-36 cells quantified per group per round. One-way ANOVA: F_(3, 8)_ = 23.32, p = 0.0003. Tukey’s multiple comparison test: **p≤0.01, ***p≤0.001. b) Rab27b KD decreases LC3II accumulation when treated with autophagosome-lysosome inhibitor chloroquine. Representative Western blot and quantification of LC3II/LC3I in isyn cells with and without Rab27b KD when induced with doxy for 48h. n = 3 independent rounds. Two-way ANOVA: CQ treatment F_(1,8)_ = 9.206, p=0.016; cell line F_(1,8)_ = 3.946, p=0.082; interaction F_(1,8)_ = 15.29, p=0.0045. Tukey’s multiple comparison test: *p≤0.05, **p≤0.01, ns = non-significant. c) Rab27b partially colocalizes with LC3 in control isyn cells upon doxy induction. Confocal image of LC3 (red) and Rab27b (green) immunoreactivity in induced isyn cells. Arrowheads point to colocalized punctae (yellow). Scale bar = 10 μm. d) Rab27b partially colocalizes with LAMP1 in control isyn cells upon doxy induction. Confocal image of LAMP1 (red) and Rab27b (green) immunoreactivity in induced isyn cells. Arrowheads point to colocalized punctae (yellow). Scale bar = 10 μm. Error bars represent SEM. D = doxy.

To further test the impact of Rab27b on autophagic flux, isyn and isyn/Rab27b KD cells were induced with doxy for 96h in serum free media followed by treatment with 40μM chloroquine for three hours. Chloroquine inhibits autophagosome/lysosome fusion, leading to an accumulation of LC3II in normal cells under autophagic-inducing conditions. Levels of LC3 II were increased in control isyn cells with chloroquine treatment, demonstrating an increase in autophagic flux (Fig. 5b). However, chloroquine failed to induce an increase in LC3 II in the presence of Rab27b KD in isyn cells (Fig. 5b). This finding points to a defect in autophagic flux with Rab27b KD. Partial colocalization of Rab27b with LC3 and with LAMP1 in isyn cells demonstrates that Rab27b can interact with autophagosomes, lysosomes, and/or autolysosomes (Fig. 5c, d). Together these data suggest that Rab27b normally promotes autophagic αsyn clearance and that impaired autophagic clearance of αsyn by Rab27b KD promotes αsyn toxicity.

### Rab27b levels are increased in human PD and DLB

We evaluated Rab27b protein expression in the post-mortem brain lysates in the medial temporal gyrus from age-matched controls and subjects with clinically and pathologically diagnosed PD or DLB. Rab27b levels were increased in the post-mortem (Triton X100-soluble) lysates of PD patients 2.1-fold compared to controls (Fig. 6a). Rab35 levels were not altered in PD brains compared to controls (Supp. Fig. 2b). Rab27b levels in the post-mortem lysates of DLB patients were also increased by 20% compared to controls (Fig. 6b).

**Figure 6.**
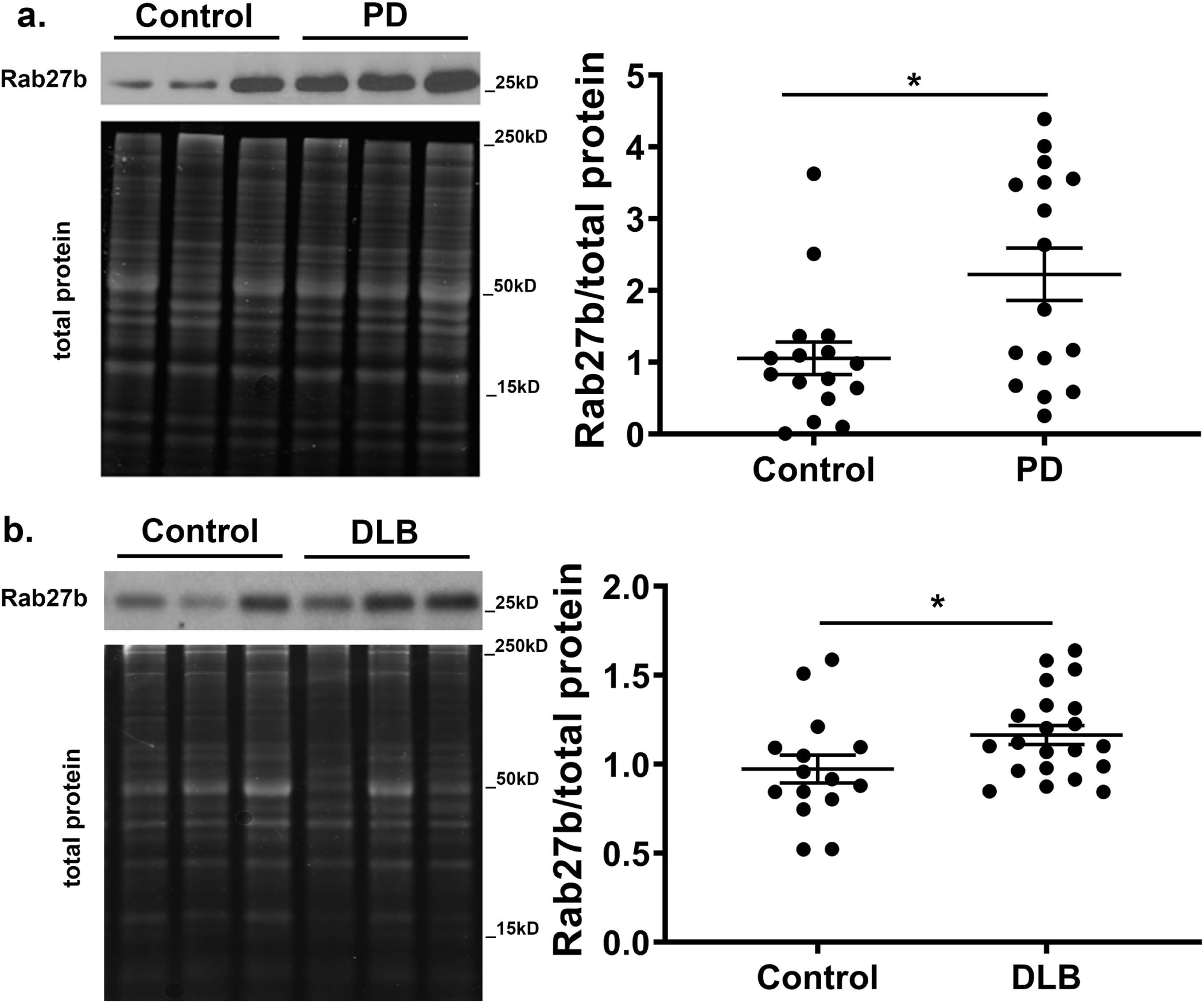
Rab27b protein expression is increased in human PD and DLB brains. a) Representative Western blot and quantification of Rab27b in control and PD post mortem temporal cortical lysates. Rab27b was normalized to total protein as determined by SYPRO Ruby Gel Protein Stain. n = 16 per group. Student’s t-test: t_(30)_ = 2.716, *p≤0.05. b) Representative Western blot and quantification of Rab27b in control and PD post mortem temporal cortical lysates. Rab27b was normalized to total protein as determined by SYPRO Ruby Gel Protein Stain. n = 15 for control and n=21 for DLB. Student’s t-test: t_(34)_ = 2.113, *p≤0.05. Error bars represent SEM.

## DISCUSSION

Our data demonstrates that Rab27b regulates the release, clearance, and toxicity of αsyn in a cellular paracrine model. We observed that Rab27b KD in isyn cells increased αsyn paracrine toxicity. Rab27b KD induced a paradoxical decrease in αsyn release, but the lower levels of released αsyn were of higher molecular weight species. Rab27b KD also increased intracellular insoluble αsyn levels. We conclude that Rab27b KD leads to an increase in αsyn paracrine toxicity due to a reduction of clearance of misfolded αsyn through autophagy. Consistent with this, Rab27b KD led to increased LC3-positive autophagosome accumulation and p62 levels and inhibited autophagic flux. Together these data suggest that Rab27b plays an integral role in the release, clearance, and toxicity of αsyn.

We have previously published on the advantages of our paracrine *in vitro* model that allows us to examine distinct parts of the various processes required for the prion-like spread of αsyn (51). The studies detailed above indicate the potential mechanisms by which Rab27b regulates αsyn spread and toxicity in this model. Our observations point to Rab27b as a regulator of αsyn propagation through multiple cellular mechanisms. Although Rab27b KD inhibited αsyn release, it also decreased autophagic flux, and released αsyn was more likely to promote toxicity. Despite the reduction in total amount of released αsyn, a probably shift to species capable of templating the misfolding of endogenous αsyn contributed to the increase in toxicity in cells treated with αsyn enriched CM from isyn/Rab27b KD cells. Our previous studies have shown that the αsyn released into the CM is primarily oligomeric and promotes seeding (51). Given that insoluble αsyn was enhanced by Rab27b KD, our data suggests that the disruption of autophagy led to an increase in oligomeric, toxic αsyn release.

Together these results indicate that Rab27b may function as an endogenous regulator of αsyn clearance via autophagy and release. Under normal conditions, we propose that Rab27b clears misfolded αsyn to prevent the intracellular buildup of toxic oligomers that can promote seeding by 1) promoting autophagic clearance and 2) promoting αsyn release (Fig. 7a). Disruption of Rab27b function allows for the accumulation of intracellular misfolded αsyn (Fig. 7b). Although total αsyn secretion is decreased by Rab27b depletion, any αsyn that is released has a higher seeding capacity (Fig. 7b). Rab27b depletion led to a decrease in αsyn release in exosomal and non-exosomal fractions, indicating that multiple release mechanisms are regulated by Rab27b and may be responsible for disease progression.

**Figure 7.**
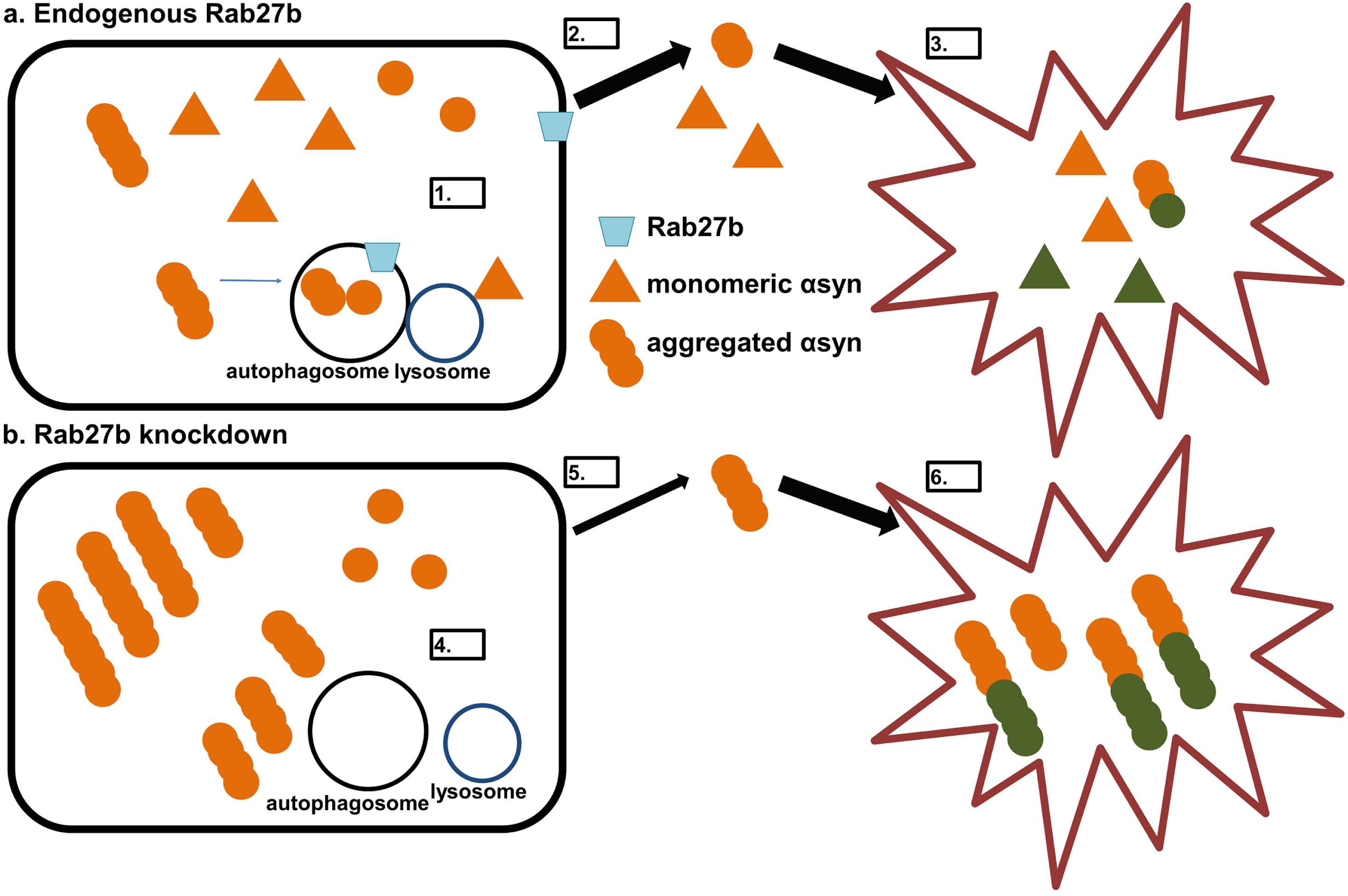
Model for Rab27b’s effects on αsyn release and clearance. a) Under normal circumstances, Rab27b decreases intracellular αsyn aggregation by increasing autophagic clearance of αsyn species (1). Rab27b also promotes the secretion of αsyn into the extracellular space (2), where the αsyn can be taken up by other neurons (3). b) Upon Rab27b knockdown, intracellular αsyn aggregation increases by impaired autophagic clearance of αsyn (4). While Rab27b KD decreases total αsyn release, the released αsyn has higher seeding potential (5), resulting in higher toxicity in neurons which take up the released αsyn (6).

Rab27b levels were increased in the post mortem brain lysates of PD patients in comparison to age-matched healthy controls. We also found that Rab27b protein levels were increased in the brain lysates of DLB patients as well, in accordance with previously published transcriptome data (50). We propose that these increases in expression may be compensatory in nature. As intracellular misfolded protein accumulates, neurons may upregulate Rab27b to increase aggregated protein clearance through the autophagic pathway. Since Rab27b also promotes αsyn release, any increase in Rab27b in disease could theoretically promote αsyn transmission from cell to cell. However, as we have previously published, an increase in total amount of αsyn released into the CM does not necessarily correlate to increased paracrine toxicity, but is instead dependent on the conformation of released αsyn (51). Indeed, our data shows that Rab27b KD actually increased the toxicity of released αsyn despite lower total αsyn amounts in the CM; increased toxicity was likely due to the release of higher molecular weight species secondary to disrupted autophagic clearance.

The molecular mechanisms by which Rab27b regulates autophagic clearance and protein secretion are unclear at this time. Rab27b has been shown to promote distal transport and docking of secretory vesicles, including lysosomes, with the plasma membrane (28,29,31,36,37,41,57-59). Rab27b could potentially promote lysosomal fusion with the plasma membrane to promote αsyn secretion or with autophagosomes to promote αsyn degradation. Rab effectors associated with Rab27b that are highly expressed in the brain include Slp5, Slp2a, rabphilin, Slac2a, and myrip (30), and we predict that different effectors are involved in the regulation of autophagy vs. release by Rab27b. Of these effectors, Slp5 (Sytl5) has been previously identified through shRNA screen by Goncalves *et al* to regulate αsyn release (18). Future directions would focus on determining the critical Rab27b effectors that regulate autophagy and secretion.

In conclusion, Rab27b regulates αsyn toxicity in our paracrine model and is upregulated in PD and DLB. Targeting Rab27b function could be a target for therapeutic intervention in these disorders.

## EXPERIMENTAL PROCEDURES

### Human brain samples

Human brain tissue was obtained from deceased persons, and the use of the human specimens was reviewed by the Institutional Review Board (IRB) at UAB and determined to be not human subjects research and not subject to FDA regulation.

### Cell lines

Isyn cells were previously created by infecting SK-N-BE(2)-M17 (M17) male neuroblastoma cells (obtained and authenticated by ATCC, Manassas, VA, Cat #CRL-2267; RRID:CVCL_0167) with the tetracycline-inducible αsyn pSLIK lentivirus in the presence of 6 μg/ml polybrene followed by selection for stable transfection with G418 (60). Isyn cells were maintained in 1:1 Eagle’s MEM/F12K containing 10% fetal bovine serum (FBS), 1% penicillin/streptomycin, and G418 (500 μg/ml) at 37°C. To induce αsyn expression, cells were treated with doxycycline (doxy) at 10 μg/ml. For Rab27b knockdown (KD) studies, isyn cells were transduced with a Rab27b targeted shRNA (5’-CCCAAATTCATCACTACAGTA-3′) (34), non-targeted shRNA (SHC016, Sigma-Aldrich), or empty vector plko.1 lentivirus, followed by selection for stable transfection with puromycin (1 μg/ml) in addition to G418 to maintain αsyn expression. Transduced cells were selected with 2 μg/ml puromycin 72h later. Lines were maintained in 1:1 Eagle’s MEM/F12K containing 10% fetal bovine serum (FBS), 1% penicillin/streptomycin, G418 (500 μg/ml), and puromycin (1 μg/ml) at 37°C. To induce αsyn expression, cells were treated with doxy at 10 μg/ml.

SH-SY5Y cells were obtained and authenticated by ATCC (Cat #CRL-2266 RRID: CVCL_0019). SH-SY5Y cells were maintained in 1:1 Eagle’s MEM/F12K containing 10% fetal bovine serum (FBS), and 1% penicillin/streptomycin. For differentiation, SH-SY5Y cells were treated with retinoic acid (10 μM) for 5-7 days in serum-free Eagle’s MEM/F12k media.

### Preparation of conditioned media (CM)

CM was prepared as previously described (51). After serial centrifugations to remove cellular debris, CM was concentrated using a 3kD Amicon Ultra-4 centrifugal filter at 4000g for two hours, followed by dialysis. Protein concentrations of CM samples were assessed by BCA assay (Thermo Fisher Scientific), and equal protein amounts were loaded for each CM sample for Western blot analysis.

For toxicity experiments, isyn cells were induced with doxy in Eagle’s MEM/F12K with 10% FBS for one week and then switched to serum-free Eagle’s MEM/F12K for 48h. Collected CM underwent centrifugation at 800g for 5 minutes, then at 2000g for 10 minutes, and then at 10000g for 30 minutes prior to transfer to differentiated SH-SY5Y cells.

### Ethidium D cell death assay

Cells were rinsed in PBS and then incubated in 1 μM Ethidium D and 2 μg/ml Hoechst 33342 in culture media for 20 minutes at 37°C. Ten high power (20X) fields per well were randomly selected for quantification, and the number of Ethidium D-positive cells and the total number of cells stained by Hoechst 33342 were counted per high power field with the rater blind to experimental conditions.

### Autophagic flux assay

Isyn cells infected with non-targeted shRNA or Rab27b shRNA were induced with doxy at 10 μg/ml for 96h in serum free EagleMEM/F12K media. Cells were then treated with vehicle or 40 μM chloroquine for three hours at 37°C prior to collection of cell lysates.

### Western blot

Western blot analysis was performed as previously described (51). Equal protein amounts were loaded per well for the CM samples and for cell lysate samples. Primary antibodies used are listed in Table 1. Blots were developed with enhanced chemiluminescence method (GE Healthcare, Piscataway, NJ). Images were scanned using the Biorad Chemidoc Imaging System and analyzed using Image Lab Biorad software for densitometric analysis of bands.

**TABLE 1.**

| PRIMARY ANTIBODIES | SOURCE | IDENTIFIER |
| --- | --- | --- |
| Mouse monoclonal anti-alpha-synuclein | BD Biosciences | Cat# 610787<br>AB 398108 |
| Rabbit polyclonal anti-alpha-synuclein | Cell Signaling Technology | Cat# 2642S<br>AB 10695412 |
| Mouse monoclonal anti-beta actin | Proteintech | Cat #66009-Ig<br>AB 2687938 |
| Mouse monoclonal anti-flotillin 1 | BD Biosciences | Cat# 610820<br>AB 398139 |
| Rabbit monoclonal anti-GAPDH | Cell Signaling Technology | Cat #2118<br>AB 561053 |
| Mouse monoclonal anti-Golgin97 | ThermoFisher Scientific | Cat #A-21270<br>AB 221447 |
| Mouse monoclonal anti-LAMP-1 | Developmental Studies Hybridoma Bank | Cat #H4A3<br>AB 2296838 |
| Mouse monoclonal anti-LC3B | Abcam | Cat #ab243506 |
| Rabbit polyclonal anti-LC3B | Cell Signaling Technology | Cat #2775<br>AB 915950 |
| Mouse monoclonal anti-p62 | Abcam | Cat #ab56416<br>AB 945626 |
| Rabbit monoclonal anti-Rab5 | Abcam | Cat #ab109534<br>AB 10865740 |
| Rabbit monoclonal anti-Rab7 | Abcam | Cat #ab137029<br>AB 2629474 |
| Mouse monoclonal anti-Rab27b | Proteintech | Cat #66944-1-Ig |
| Mouse polyclonal anti-Rab27b | Abcam | Cat #ab76779<br>AB 1524280 |
| Rabbit polyclonal anti-Rab27b | Millipore-Sigma | Cat #abs1026 |
| Mouse monoclonal anti-α-tubulin | Sigma-Aldrich | Cat #T9026<br>AB 477593 |

Fresh-frozen tissue from temporal cortices of age and gender-matched control, PD, and DLB brains were obtained from Banner Sun Health Research Institute Brain and Body Donation Program. Samples were prepared as previously described (61). Rab27b protein levels were normalized to total protein levels determined by SYPRO Ruby protein gel stain (Invitrogen).

### Exosome fractionation

For exosome preparation, we followed the protocol described in (51,54). Briefly, cells were incubated to serum-free Eagle’s MEM/F12K for 96h. CM underwent serial centrifugations at 800g for 5 minutes, 2000g for 10 minutes, and 10000g for 30 minutes to remove cellular debris at 4°C. Debris-free media was then spun at 100000g for two hours at 4°C. The supernatant was saved as the non-exosomal fraction. The pellet was resuspended in PBS and then spun again at 100000g for 80 minutes at 4°C. The pellet was resuspended in PBS with protease inhibitors.

### Nanosight

The size of exosomes was determined on a NanoSight NS300 (Malvern Instruments, Westboro, MA), as previously described (51). Following isolation by ultracentrifugation, exosome pellets were resuspended in 60μl cold PBS with repeated pipetting and vortexed for 10 seconds. Then 15μl of the exosome suspension was diluted to a total volume of 1ml PBS and analyzed on a NanoSight NS300 infused with a syringe pump set at 25 (arbitrary units). Data were collected for each sample in 10 repeats of 60 second video and analyzed using NanoSight NTA 3.0 software.

### Size exclusion chromatography

Size exclusion chromatography (SEC) was performed as previously described (51). 20 μl (80 μg) of CM diluted in PBS was loaded onto a NGC FPLC (Bio Rad Laboratories), injected on a Yarra 3 μm SEC – 2000 column (300 x 7.8 mm, Phenomenex), and run at 0.7 ml/minute in 1x PBS, pH 6.8. 250 μl fractions were collected from elution volume 4 – 12.5 ml. This corresponds to the end of the void volume, as determined by a Blue Dextran standard, and the buffer front, as determined by imidazole elution. αSyn in 50 μl of each fraction was measured using an ELISA for αsyn.

### Immunocytochemistry

Isyn cells were fixed in 4% paraformaldehyde for 15 minutes. After washing in PBS, cells were permeabilized with 0.5% Triton X-100 in PBS for 20 minutes and then blocked with 5% NGS in PBS for 20 minutes. Cells were incubated overnight with primary antibody (Rab27b, Rab5, Rab7, LC3II, p62, Golgin97, LAMP1) in 1.5% NGS. Primary antibodies used are described in Table 1. After washing, cells were incubated with goat anti-rabbit or anti-mouse secondary antibody in 1.5% NGS for 2h. Isyn cells were imaged using an Olympus BX51 epifluorescence microscope. Ten high power (20X) fields per well were randomly selected for quantification, and the immunoreactivity was quantitated using ImageJ with the rater blind to experimental conditions. For LC3II puncta counts, slides were imaged at 63x by confocal (Leica TCS-SP5 Laser scanning confocal microscope) and quantitated using ImageJ cell counter. For Rab27b colocalization, Z-stack images of neurons were taken at 63x by confocal (Nikon Eclipse Ti2 scanning confocal microscope).

### Experimental design and statistical analysis

GraphPad Prism 8 (La Jolla, CA) was used for statistical analysis of experiments. Data were analyzed by either student t-test or by one-way or two-way ANOVA, followed by post-hoc pairwise comparisons using Sidak’s or Tukey’s multiple comparison tests. Statistical significance was set at p ≤ 0.05. ANOVA related statistics (F statistic, p values) and post-hoc test results are found in the figure legends. For t-tests, the t statistic and p values are noted in the figure legends.

## ACKNOWLEDGEMENTS

We are grateful for Drs. Geidy Serrano and Thomas Beach at the Banner Sun Health Research Institute Brain and Body Donation Program of Sun City, Arizona for the provision of human brain tissue. The Brain and Body Donation Program is supported by the National Institute on Aging (P30 AG19610 Arizona Alzheimer’s Disease Core Center), the Arizona Department of Health Services (contract 211002, Arizona Alzheimer’s Research Center), the Arizona Biomedical Research Commission (contracts 4001, 0011, 05-901 and 1001 to the Arizona Parkinson’s Disease Consortium) and the Prescott Family Initiative of the Michael J. Fox Foundation for Parkinson’s Research. Research reported in this publication was also supported by the UAB High Resolution Imaging Facility.

This study was supported by NIH [R01 NS088533 (TAY); R01 NS112203 (TAY); P50 NS108675 (TAY); R01GM117391 (WJP); F31 NS106733 (RU)], American Parkinson Disease Association, and the Parkinson Association of Alabama. The content is solely the responsibility of the authors and does not necessarily represent the official views of the National Institutes of Health.

## AUTHOR CONTRIBUTIONS

RU designed and performed experiments, analyzed data, and wrote manuscript. BW designed and performed experiments, and reviewed manuscript. RWH performed experiments and analyzed data. WJP, RWH, and CC designed experiments and reviewed manuscript. TY designed experiments, analyzed data, and edited and finalized manuscript draft.

## List of abbreviations

αsyn: alpha-synuclein
CM: conditioned media
DLB: Dementia with Lewy Bodies
doxy: doxycycline
EV: empty vector
isyn: doxycycline-inducible alpha-synuclein cell line
KD: knockdown
KO: knockout
PD: Parkinson’s disease

## FIGURE LEGENDS

**Supplemental Figure 1:**
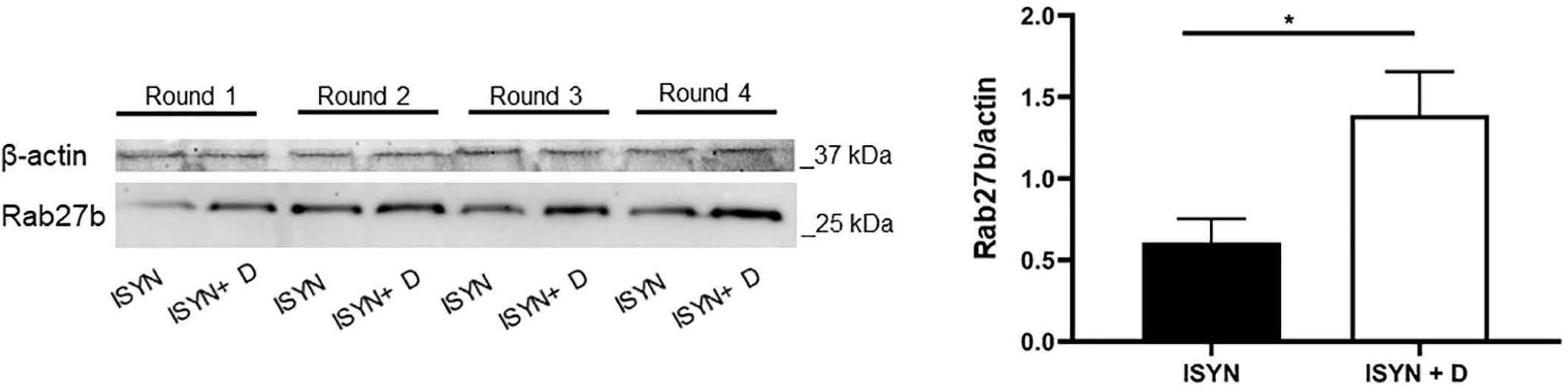
Rab27b levels are increased in an αsyn overexpression model. Representative Western blot and quantification of Rab27b protein levels upon αsyn overexpression for 96h. n = 4 independent rounds.

**Supplemental Figure 2.**
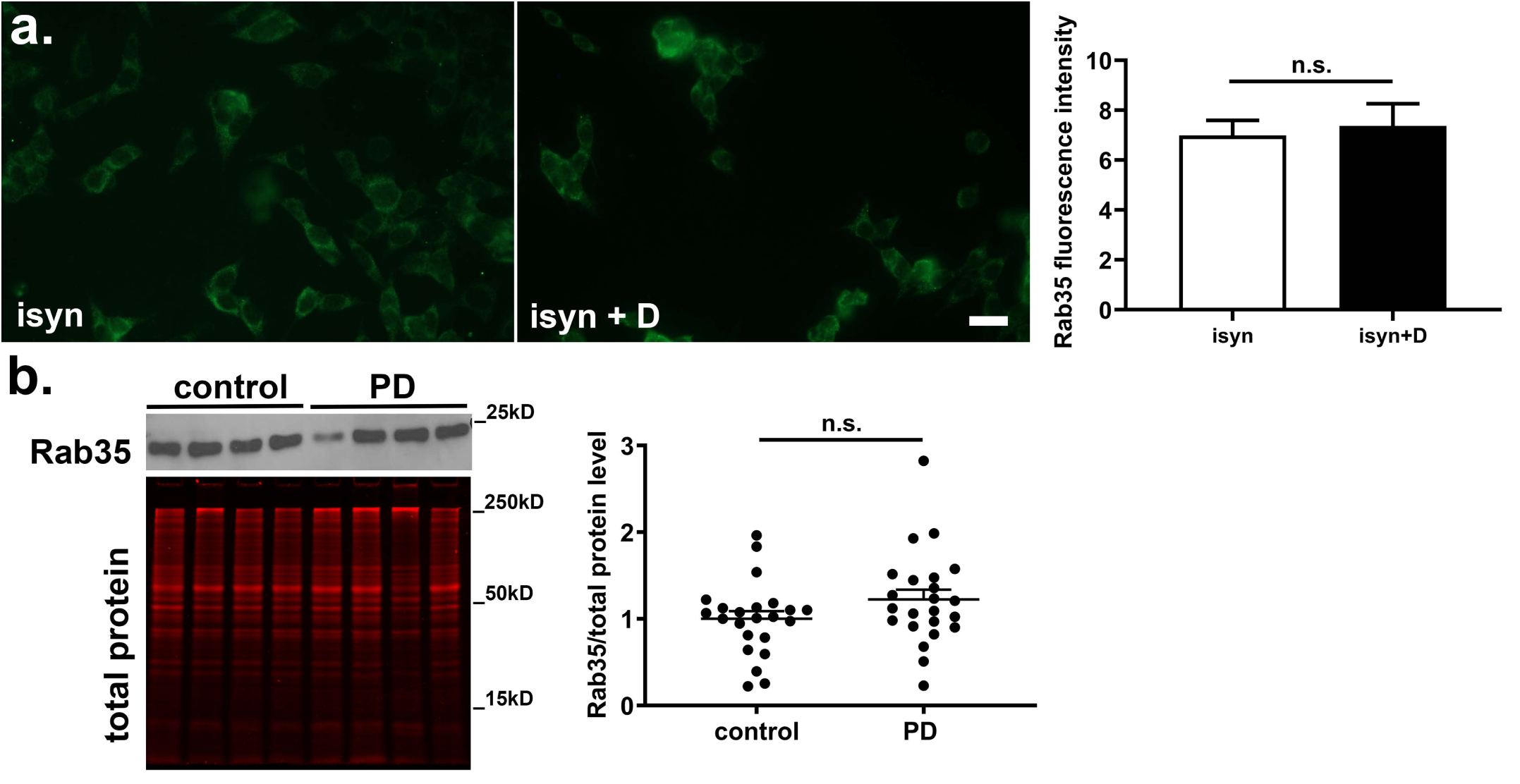
Rab35 levels are not altered in a paracrine αsyn model or in human PD. a. Rab35 immunoreactivity in isyn cells upon αsyn overexpression for 48 hours. n= 3 independent rounds. b. Representative Western blot and quantification of Rab35 in control and PD post mortem temporal cortical lysates. n= 23/group.

